# Cryptic and Ecologically Relevant Locomotor Activity Patterns in Lake Malawi Cichlids Revealed Through Computer Vision Pipeline

**DOI:** 10.1101/2025.05.15.654275

**Authors:** Niah Holtz, Evan Lloyd, Chloe Hoff, Alex C. Keene, R. Craig Albertson

## Abstract

Cichlid fishes have long been a popular model for ecology and evolutionary biology. Not only do they exhibit nearly unparalleled taxonomic diversity, but their community structures are both complex and extremely dense. Niche partitioning along dietary axes has been credited in maintaining cichlid biodiversity in a broad sense; however, it is common for cichlid species with overlapping diets to co-exist, leading to the search for other mechanisms to explain the maintenance of cichlid biodiversity. Behavioral variation has the potential to lead to fine-scale (i.e., micro-) habitat partitioning, but analyzing complex animal behavior poses several challenges, including time, cost and reproducibility. Modern tools, such as Machine Learning (ML), ease the burden of behavioral analyses, and are increasingly used in the field. Here, we present the application of tools from a well-developed sub-field of ML, Computer Vision (CV), to extract locomotor behavioral data from four Lake Malawi cichlid species. Our dataset consisted of low-resolution (320×240), twenty-four-hour infrared videos of activity in 4 cichlid species. We first analyzed a pair of species with divergent behaviors, diets and habitats (i.e., *Abactochromis labrosus* versus *Tropheops kumwera*), representing a proof-of-concept comparison. Next we analyzed activity in two closely related species with broadly similar behaviors and overlapping diets and habitats (i.e., *Maylandia* sp. ‘Daktari’ versus *Maylandia faiaziberi*). In both comparisons, our analysis involved the quantification of tank usage patterns and stop/rest activity. Not only did our data align with the known ecologies and behaviors of the species in our proof-of-concept comparison, but they revealed novel behavioral differences between all species, especially as related to variation in tank usage from day to night. Our approach provides a window into potential cryptic behavioral niche partitioning, as well as a foundation for future, high-throughput analyses to study the genetic basis and evolution of behavior.

## INTRODUCTION

Animal behavior is complex and dynamic and has increasingly become a field that relies on the rigorous and reproducible quantification of large datasets. The time-consuming nature of manual quantification can limit the pace of progress, and associated issues with reproducibility can confound behavioral experimentation (Baker 2016; Branch 2018). The application of machine learning (ML) and other automated methodologies has the potential to transform the analysis of animal behavior by allowing investigators to ask more complex questions, analyze larger datasets, and reveal cryptic patterns.

Computer Vision (CV) is a field focused on applying ML models to visual data, allowing computers to interpret and classify visual information (Wiley and Lucas, 2018). Given its ability to detect patterns from complex datasets, CV has become increasingly popular in the field of animal behavior (Ardekani, 2013; Palmér, 2017; Graving, 2021). For example, CV has been used to detect and classify animals in camera trap images (Schneider et al., 2018; Ferrante et al., 2021), providing a boon to animal conservationists and invasive species specialists alike. It has also been widely used to observe stress behaviors and the wellness of livestock, such as pigs, chickens, and fishes (Leroy et al., 2005; 2006; Papadakis et al., 2012; Lao et al., 2012; Wurtz et al., 2019; Fernandes et al., 2020; Liu et al., 2020; Chen et al., 2021; Li et al., 2021; Borges Oliveira et al., 2021; Bhuiyan et al., 2023), alongside domesticated pets, such as dogs (Barnard et al., 2016; Chávez-Guerrero et al., 2022). Moreover, CV has been used to classify animal behaviors in the wild, including differentiating between different postures in foxes (Schütz et al., 2022), as well as collective measurements of sociality and migration in populations of storks, ungulates, and primates (Flack et al., 2018; Walter, 2022; Koger et al., 2023). CV also holds great potential to understand animals’ umwelt better, offering new insights into long-standing questions concerning how animals perceive the world (Okuyama et al., 2015; Mathis and Mathis, 2020; Broomé et al., 2022; Banerjee et al., 2021; Vogg et al., 2024). That perception, alongside a complex interplay between behavior, life history, and ecology, ultimately influences an animal’s survival and reproductive success. With a rapidly expanding CV toolkit, we are gaining deeper insights into how and why certain species are ecologically and evolutionarily successful.

Cichlids are a family of fishes that have long been a popular model for ecology and evolutionary biology owing to the speed of their diversification and their complex community structures (Kornfield and Smith, 2000; Seehausen, 2006; Gavrilets et al., 2007; Arbour and López-Fernández, 2014; Burress, 2015; Koblmüller et al., 2019; Jordan et al., 2021). They first appeared in the fossil record 46-55 mya before the breakup of Gondwana (Genner et al., 2007). Today cichlids thrive as one of the largest vertebrate families with nearly two thousand formally described species and an actual number that is likely much higher (Kullander, 1998; Santos et al., 2023). While they exhibit a circumtropical distribution, cichlid biodiversity is predominantly centered within freshwater rivers and lakes in South America and Africa, where their evolutionary success is credited to extensive and iterative bouts of adaptive radiations (Kocher, 2004). Equally impressive is the high density of cichlid species in their natural environment. For instance, within Lake Malawi, it is normal for up to a dozen species with overlapping foraging niches to be present at a particular location (Ribbink et al., 1983). Such observations have led investigators to search for axes of habitat partitioning to explain how these complex species communities are maintained.

Niche partitioning in cichlids is well studied along dietary and spatial axes. For example, isotope analyses support the existence of dietary niches (Bootsma et al., 1996; Genner et al., 1999; Genner et al., 2001), but these methods cannot distinguish between what an animal eats and how it gathered that prey item. Phytoplankton, for example, may be grazed from rocks upon which it settles or collected directly from the water column, conflating dietary and spatial habitat similarity. The concept of micro-habitats (Gould, 1837; MacArthur, 1958) suggests that species may share a dietary niche provided food is gathered at different locations and/or in a different manner. This concept implies that close ecological competitors should evolve different foraging strategies and morphologies (Adams and Huntingford 2002; De León et al., 2014; Skoglunch et al., 2015; Ingram et al., 2022). A thin, narrow jaw, for example, can generate suction to harvest plankton from the water column, while a wide, stout jaw is better suited for scraping the same plankton that has settled onto rocks. Indeed, cichlid foraging architecture is an accurate predictor of not just what the species eats but how food is gathered (Cooper et al., 2010; Sampaio et al., 2013; Martinez et al., 2018), and even subtle differences in foraging habitat are reflected in the shape of the cichlid feeding apparatus (Albertson and Pauers, 2019; Conith et al., 2020). Cichlids are also well known for their ability to adapt plastically to changes in foraging opportunities (Parsons et al., 2016; Carruthers et al., 2022; Tetrault et al., 2024). Such behavioral polymorphism allows organisms to respond to the environment, specializing or generalizing their foraging behavior as opportunity allows (Parsons et al., 2010; Doenz et al., 2019; Fenton et al., 2024). In other words, the minutiae of fish behavior can snowball into long-term ecological and evolutionary trends.

While the questions of “where” and “how” have been well studied in the realm of cichlid niche partitioning, the question of “when” is generally lacking. Time can be an important factor in determining habitat/resource availability, and includes seasonal and circadian variability (Martin and Genner, 2009; Soria-Barreto et al., 2008; Nichols et al., 2024). Historically, circadian patterns of behavior have gone largely unstudied in cichlids; however, recent work suggests that this is a fruitful area of study. Lloyd and colleagues, for example, documented day-night variation across Lake Malawi species, including strongly diurnal, strongly nocturnal, and arrhythmic patterns (Lloyd et al., 2021). In addition, Nichols and colleagues surveyed day-night activity in 60 Lake Tanganyikan cichlids and showed that species with similar diets tended to exhibit divergent chronotypes, consistent with temporal niche partitioning (Nichols et al., 2024). We seek to extend these studies by applying ML to quantify variation in tank usage patterns and timing among four exemplar Lake Malawi cichlids. In order to investigate how niche partition and activity patterning evolved in cichlid fish, we sought to develop methodologies for high-throughput, automated analyses of locomotor activity over twenty-four-hour periods. Specifically, we used multiple CV tools to quantify multiple spatio-temporal aspects of locomotor activity. Our analysis provides a mainly open-source system for long-term behavioral measurements in cichlids that can readily be applied to other fish species.

## MATERIALS AND METHODS

### Fish species and husbandry

Cichlid species in this study include *Abactochromus labrosus*, *Tropheops kumwera*, *Maylandia sp*. “Daktari”, and *Maylandia faiaziberi*. Fishes were obtained as juveniles through the aquarium trade and reared according to standard protocols approved by the IACUC from UMASS Amherst and TAMU fish facilities. Water temperatures were 28.5°C, and lighting was on a 14:10 schedule from light to dark. Fishes were reared in conspecific groups in 40 gallon glass aquaria, and fed a high quality flake food mixture of ∼2/3 spirulina algae and ∼1/3 yolk flake once daily. Groups of 5 experimental fish (n=2 males, n=3 females) were moved to 10 gallon glass filming tanks for a twenty-four-hour acclimation period before filming. Animals were fed and the water was changed between the acclimation and filming periods.

### General CV terminology and techniques

The power of CV tools is growing every day, but the application of these tools can be overwhelming due to their complexity and associated jargon. As with most CV approaches, we use a Convolutional Neural Network (CNN) to extract ‘features’ of ‘classes’ from a labeled subset of representative images, in order to ‘train’ our ML model to ‘detect’ these features in a larger dataset. Each of these steps or categories has a plethora of considerations to ensure the appropriate and efficient use of the tool at hand. For example, the underlying architecture of any given CNN is highly varied based on the task it was trained to do and the resources available (Zhao et al., 2024). Further, several mathematical layers process standalone images or video frames to determine the features uniquely associated with labeled areas of interest. For those interested in the minutiae of this process, we advise getting started with the publicly available Stanford course on Convolutional Neural Networks for Visual Recognition (Jiang, 2015).

Features depend on the model’s training and, thus, the capability to quantify the data’s composition, context, and type of labeling (e.g., centroid versus bounding box). For example, when seeking to quantify how many times a green insect lands on a green plant, green—or its numerical representation—might not serve as a useful feature. Moreover, postural features will be indeterminable in data labeled solely with a bounding box. Common features include, but are not limited to, edges, textures, colors, points or corners that contrast with the background (Selfridge, 1959; Lindeberg, 1998; Lowe, 2004). Once features are identified, typically by training a relevant existing model on the labeled data, they can associate with classes in downstream detections. A class is programmatically established during the initial labeling process prior to training and represents the object of interest (Deng et al., 2010; Parisot et al., 2023). Like in the example above, a class would be the insect itself labeled in each photo and mined for unique features during training. Once classes are established, the data are labeled, and a model is chosen; then, training begins with a labeled dataset split into two parts: training and testing. In this way, additional labeling only occurs if extra training is needed, which depends on the number of images labeled to start and the complexity of the dataset compared to the data on which the baseline model was trained. This process can and should occur iteratively as it becomes apparent how the model is performing in detecting the defined class.

Model performance can be checked through statistical measurements and practical application. Widely accepted metrics start with precision (i.e., true detections over total detections) and recall (i.e., true detections over total true objects). There is an inherent tradeoff between the two; if missing a detection is preferred to having a false detection, one would prioritize high precision and vice versa (Oyen, 2013). If neither is appropriate, one optimizes for mean Average Precision (mAP), a curve based on the precision and recall of a particular dataset alongside two other metrics, the Intersection over Union (IoU) and Confusion Matrix (Henderson and Ferrari, 2016; Dave et al., 2022). Higher IoU shows better alignment between actual labels and model predictions. The Confusion Matrix compares each actual detection with model predictions. These metrics assess model performance, but testing on unlabeled data reveals the strengths and weaknesses of the training, a powerful way to quickly generate new labeled data to train a model on features that may be poorly represented. In our case, the model was reluctant to identify a fish swimming straight upward, so we extracted many images of this occurring and trained specifically on those to improve model performance. Once satisfied with what can be seen by running the model on random samplings of the dataset and assessing statistical metrics, one can begin generalizing these predictions to the rest of the dataset. All CV applications must rely on a ‘trust but verify’ ideology; with that said, we hope this section helps explicate some of the jargon associated to CV tool usage

### Data preparation

We prepared our dataset for training through trainYOLO (Redmon et al., 2016) using FFMPEG (FFMPEG Developers, 2016), OpenCV (Culjak et al., 2012), and an augmented DEEPLABCUT script (Mathis et al., 2018). Our dataset consisted of twenty-four-hour videos (320×240) of four species of Lake Malawi cichlids (*Abactochromis labrosus, Tropheops kumwera, Maylandia* sp. ‘Daktari’, and *Maylandia faiaziberi*), obtained using an augmented Microsoft Lifecam with an Edmund Optics IR long-pass filter through VirtualDub2 (Figure 1a). The videos captured five conspecific fish in glass aquaria illuminated by LED IR 850 nm lighting strips and lined with white corrugated plastic sheets (Figure 1b). Our CV pipeline is represented as a flowchart in Figure 1c and described below. We used FFMPEG to crop each tank and trim the videos to approximately four hours for optimal computation during tracking. Then, a random subset of these videos was streamed using OpenCV, and frames were selected using MiniBatchKMeans from sklearn.cluster, returning a user-provided number of frames from associated visual clusters (Pedregosa et al., 2018). Extracted frames were then uploaded to trainYOLO for easy annotation compatible with our model. Open-source alternatives to this labeling tool include, but are not limited to, LabelStudio (Tkachenko, 2020), COCO Annotator (Stefanics and Fox, 2022), and Imglab (Trejo and Angulo, 2016). Training via the command line is also possible. After training, we downloaded our model weights for inference and subsequent tracking of video data.

**Figure 1.**
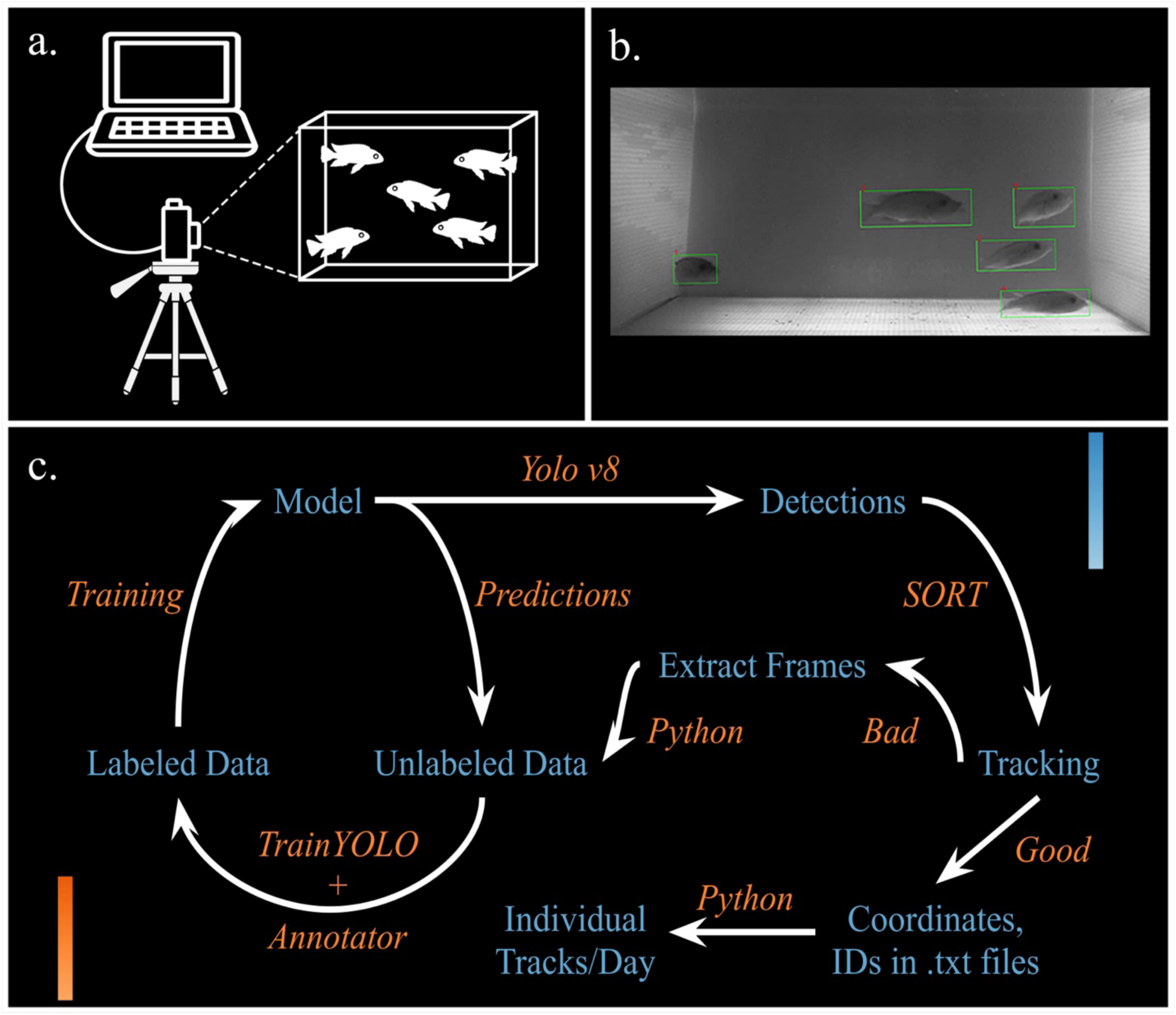
Graphical Methods of Filming and Data Analysis. a) Visual representation of experimental setup, five fish in a single tank filmed via augmented Microsoft Lifecam. b) Labeled frame from a twenty-four-hour film. c) Experimental pipeline flowchart, start with unlabeled data. The blue text refers to the data being manipulated, while the orange text refers to the tools used to manipulate the data.

### Inference and tracking

Inference and tracking were performed using OpenCV, YOLOv8, and an augmented SORT script (Bewley et al., 2017). We analyzed the prepared videos en masse by streaming each frame individually with OpenCV and conducting inference on each frame using the yolov8n model to return detections with a confidence of 0.5 and an IoU of 0.2 for our selected class. These detections were passed to our augmented SORT script. SORT uses a Kalman filter to generate tracks and match future detections to tracks. Here, user augmentations limit the track number by the known number of animals (5 in our case). Forcing SORT to choose the ‘best’ match of all available tracks for each detection rather than create new tracks needing future stitching. As with all Multi-Object Tracking (MOT) methods, the procedure is vulnerable to identity switching (Yilmaz et al., 2006). Thus, our interpretation of the tracked data was limited to group (not individual) dynamics. The final tracks were returned to the original script for storage by pickling, a type of data serialization that reduces stored file size, and the process repeats until each cut from a single original video is processed. At that point, the KalmanBoxTracker.count was set to zero, and all pickled outputs were grouped for further analysis and quality checks.

### Quality control

Quality checks occurred during tracking, after pickle output, and before final visualizations through matplotlib.pyplot. We used OpenCV for live video tracking, manually checking the stream for issues. After monitoring, file size was inspected to ensure it fell within a reasonable range of the median size of all videos. If too small, this likely meant that the model was unable to detect the fish in a significant number of frames, so we undertook additional training using targeted frames. If too large, videos were reviewed for the presence of objects that could be confused for a fish, an easy mistake to make in low-resolution IR video. Videos that failed these standards after two rounds of training and re-tracking were disqualified.

### Data visualization and analysis

Given a pass in all stages of quality checks, data moved forward for visualization through heat maps of percent-of-total presence and scatterplots of stopping behavior. Heatmaps binned activity into 6×3 spatial bins. Day, night, and day-night heatmaps were made from coordinate data and included the percent presence in each bin. Scales specific to each graph are presented to the right, with darker colors indicating higher percentages. We used variance of presence to approximate spatial preference in day and night heatmaps, such that high variance indicated a high preference for a particular region of the tank, whereas low variance indicated low spatial preference. For day minus night heatmaps, variance approximates shifts in spatial preference; high values indicate a strong shift in spatial preference between day and night, whereas low indicate no or weak shifts. For these graphs, the scales indicate the difference in percentage between day and night, with positive values indicating greater time spent in a bin during the day and negative values indicating more time at night.

Since the percent time spent in an area of the tank could mean that fish were either stopped in that location or swimming back and forth in a limited area, we next generated scatter plots for day and night based on where and the amount of time a fish stopped. For this analysis, a “stop” indicated that an animal did not move more than one body length in any direction for sixty seconds or longer. The location of each dot indicates where a stop was detected, and the size and color indicate the length of a stop. A darker color indicates a longer stop relative to all other stops within the video. The size of the dot indicates the absolute time of the stop. Each plot also included the percent time spent stopped, as well as the standard deviation and error of that measurement. The scales at the right for these graphs show the range of time (in seconds) for all recorded stops.

## RESULTS

### Behavioral patterns align with known ecological niches

Our initial comparison was between two heterogenetic species, *Abactochromis labrosus* and *Tropheops kumwera*. *A. labrosus* is a weakly territorial species with a proclivity for crevices and benthic crustaceans (Figure 2a). Alternatively, *T. kumwera* is highly territorial and spends much of its time above rocks, feeding on algae or interacting with other cichlids (Figure 2b); our previous work has documented high levels of activity for this species (Lloyd et al., 2021; 2024). From this information, we expected *A. labrosus* to have a higher spatial preference than *T. kumwera*; a prediction that was born out by our data. Specifically, *A. labrosus* exhibited a strong spatial preference for the top left corner during the day (e.g., 69% of the time; Figure 2c). At night, *A. labrosus* exhibited a similar pattern, though it was not as strong, with animals also spending time along the bottom and sides of the tank (Figure 2e). This pattern was supported by relatively high variances for both day and night usage, 245.05 and 15.07, respectively. All in all, this species strongly preferred the corners, sides, and bottom of the tank, actively avoiding the middle. *T. kumwera* exhibited a very different pattern, with day and night heatmaps showing relatively even presence across all spatial bins and correspondingly low variance scores of 0.95 and 2.13 for day and night, respectively (Figure 2d, f). The pattern was slightly weaker at night, with a slight preference for the bottom of the tank.

**Figure 2.**
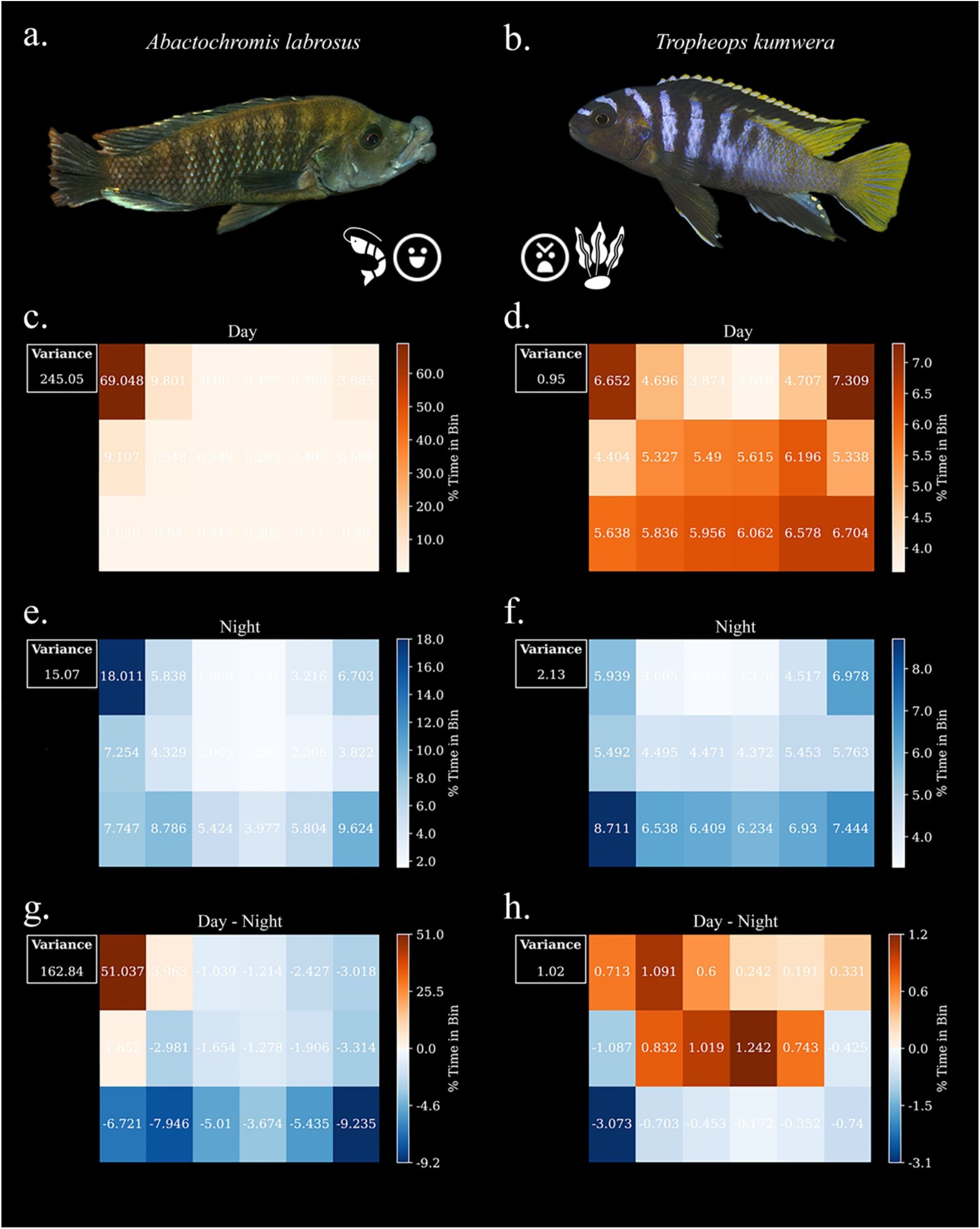
Heatmaps of Fish Tank Usage (*Abactochromis labrosus* and *Tropheops kumwera*). Photos of each species courtesy of Ad Konings and Cichlid Press. Pictured left (a), *A. labrosus*, a relatively non-territorial species with a largely crustacean-based diet. Pictured right (b), *T. kumwera*, a highly territorial herbivorous species. Heatmaps to the left (c, e, g) represent *A. labrosus,* and those to the right (d, f, h) represent *T. kumwera.* To the left of each heatmap is the variance of time spent in each spatial bin. To the right is a gradient scale depicting a range in the percent time spent in each spatial bin. Note that the scale range can differ markedly between plots, and pattern variables are more informative when coinciding with high variance. Data are presented as day (c-d) and night (e-f) heatmaps, showing the pattern of presence related to time with darker colors correlating with higher presence. Day minus night heatmaps (g-h) highlight the change in usage patterns from day to night, with darker reds correlating with higher presence during the day and darker blues correlating with higher presence during the night. High variance values in these plots may be interpreted as larger differences between day and night usage patterns.

Day minus night heatmaps and variances confirmed a strong difference in tank usage for *A. labrosus*, with a variance of 162.84 and a clear preference for the upper left region of the tank during the day, and a more free roaming pattern at night (Figure 2g). *T. kumwera* had a much lower variance of 1.02, indicating little difference in spatial preference between day and night (Figure 2h). The difference between these two species was further underscored when comparing the range of percentages, which spanned −9.23 to 51.04 for *A. labrosus* but only −3.073 to 1.242 for *T. kumwera.* These data stand as a good proof-of-concept as they align with what is known about these species with divergent ecological niches; however, they also reveal previously undescribed behaviors for *A. labrosus*, particularly in terms of tank usage at day versus night.

### Pattern segregation in closely related species with shared ecological niches

We next sought to compare two closely related species, *Maylandia sp.* “Daktari” and *Maylandia faiaziberi,* that share dietary proclivities as algalvores and tend to spend time above the rocky substrate as adults (Figure 3a-b). They have overlapping distributions in Lake Malawi and are weakly territorial (Cicotto et al., 2011). Thus, we had no *a priori* prediction as to how or if these species would segregate behaviors.

**Figure 3.**
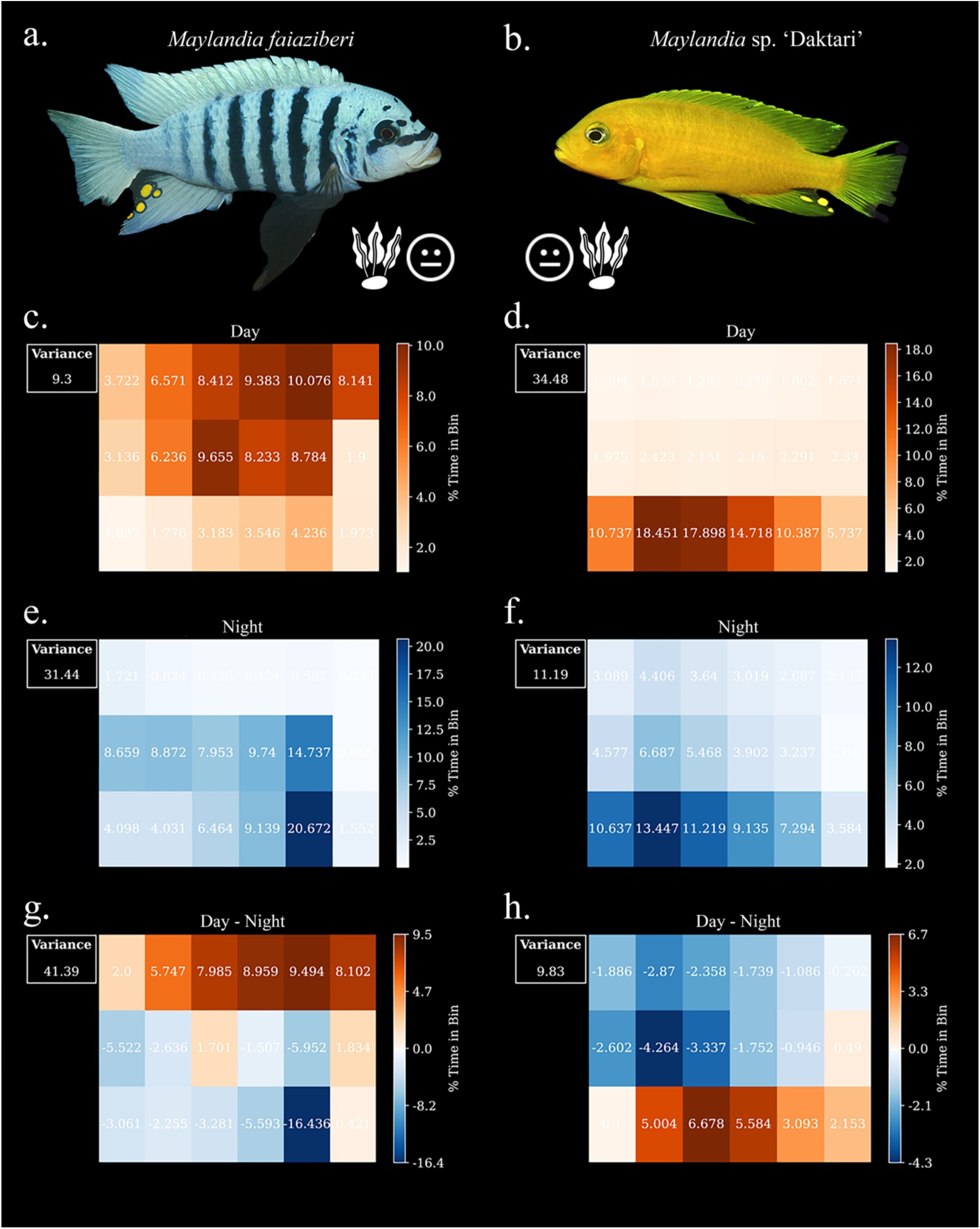
Heatmaps of Fish Presence (*Maylandia faiaziberi* and *Maylandia sp*. ‘Daktari’). Photos of each species courtesy of Ad Konings and Cichlid Press. Pictured left (a), *M. faiaziberi*, a weakly territorial herbivorous species. Pictured right (b), *M. sp*. ‘Daktari’, also a weakly territorial herbivore. Heatmaps to the left represent *M. faiaziberi,* and those to the right represent *M. sp*. ‘Daktari’. To the left of each heatmap is the variance of time spent in each spatial bin. To the right is a gradient scale depicting a range in the percent time spent in each spatial bin. Note that the scale range can differ markedly between plots, and pattern variables are more informative when coinciding with high variance. Day (c-d) and night (e-f) heatmaps show the usage pattern as related to time, with darker colors correlating with higher presence. Day minus night heatmaps (g-h) highlight the change in usage from day to night, with darker reds correlating with higher presence during the day and darker blues correlating with higher presence during the night. Variance values in these plots relate to differences between day and night usage patterns.

Both day and night heatmaps for *M. sp.* “Daktari” showed a decided preference for the bottom of the tank with higher variance during the day (34.48) than at night (11.19) (Figure 3c, e). Notably, *M. faiaziberi* exhibited a different pattern with a higher preference for the middle/top of the tank during the day (variance = 9.3) and for the middle/bottom at night (variance = 31.44) (Figure 3d, f). This difference is reflected in the day-night heatmaps, where *M. sp.* “Daktari” had a slight day-time preference for the bottom of the tank (variance = 9.83) and *M. faiaziberi* exhibited a stronger day-time preference for the top of the tank (variance = 41.39) (Figure 3g-h).

### Stopping behaviors underlie patterns of tank usage

We next used stop detection to determine the specific behaviors that underlie our tank usage patterns. For example, it is possible that preference for a particular area of the tank, as determined by our heat maps, was due to constant swimming within that area (e.g., along the bottom of the tank) or due to inactivity within that area. Stop detection data incorporates time into presence/absence data, thereby discriminating between stationary and mobile behaviors within a particular area. *A. labrosus* exhibited the highest spatial preference of the four species examined here, especially during the day, and we noted a correspondingly high level of stopping in the same area. In fact, *A. labrosus* spent 82.5% of the time stopped during the day (Figure 4a), whereas at night this species was only stopped 9.6% of the time. This pattern strongly suggests a nocturnal pattern of rest/activity, something noted for other Lake Malawi cichlids (Llyod et al., 2021; 2023) (Figure 4b). In contrast, *T. kumwera* stopped very little overall, 0.29% during the day and 1.02% at night (Figure 4c-d). Moreover, these stops occurred throughout the tank, consistent with the tank usage data.

**Figure 4.**
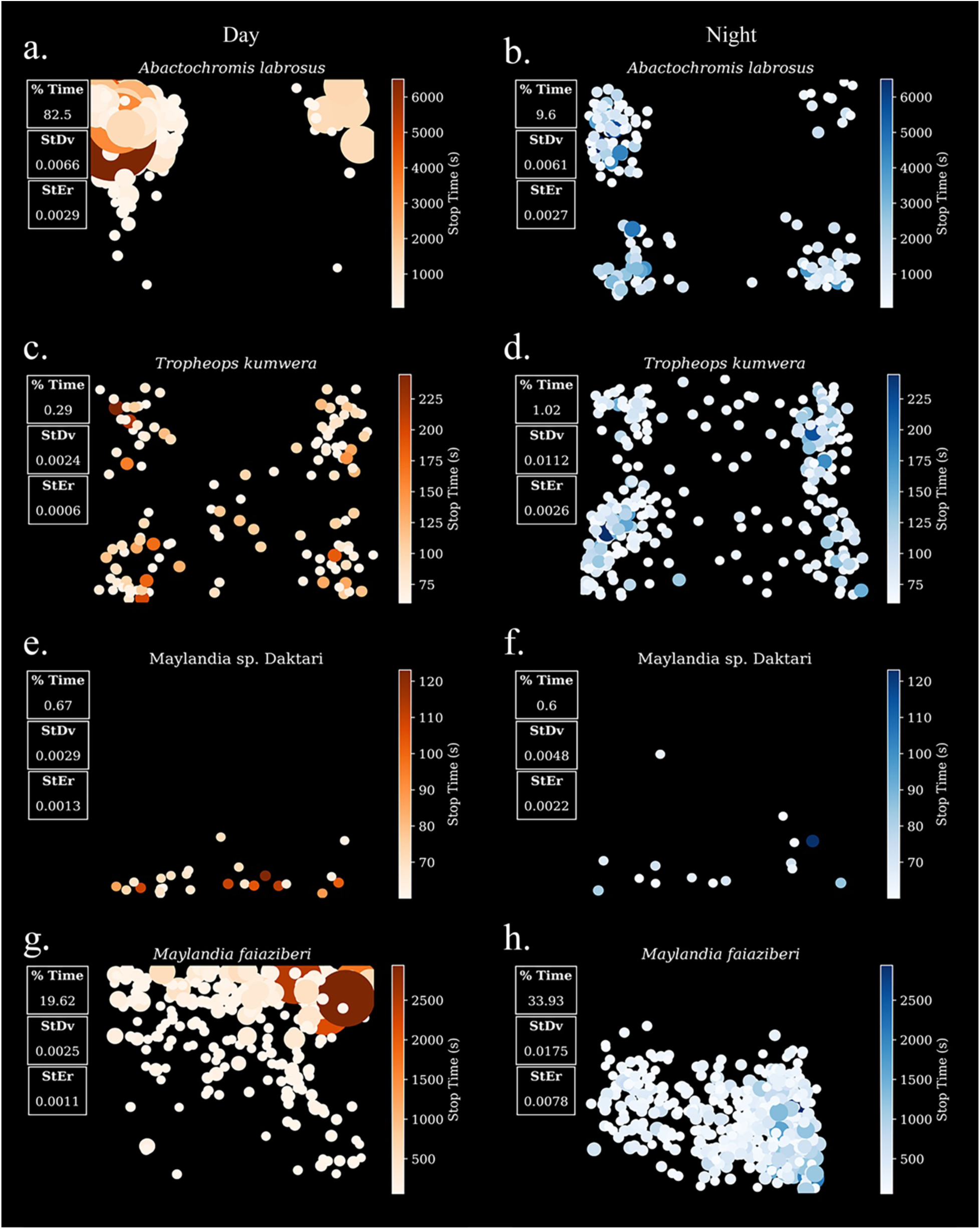
Stop Detection Data. Scatterplots to the left represent day stops (orange), while those to the right indicate stops at night (blue). Each row represents one of the four species analyzed: *Abactochromis labrosus (a-b), Tropheops kumwera (c-d), Maylandia sp*. ‘Daktari’ (e-f), and *Maylandia faiaziberi* (g-h). Each dot represents the location of one stop for 60 consecutive seconds or more; the longer the stop, the larger the dot. Since dot size represents absolute time, it may be compared across all plots. To the left of each scatterplot is the percent time spent stopped during the day (a, c, e, g) and night (b, d, f, h) alongside the standard deviations and errors of the duration of each stop. To the right of each scatterplot is a gradient scale of time spent stopped in seconds; the darker the color, the longer the stop relative to other recorded stops in the same plot. Note that the scales are very different for each species, and thus dot color is only relevant within a single plot.

For the congeneric comparison, tank usage data showed that *M. sp.* “Daktari” preferred the bottom of the tank during the day, while *M. faiaziberi* preferred the top. Our stop data provided additional insights into how each species used those areas. Specifically, *M. sp.* “Daktari” stopped very little and for very short periods of time at the bottom of the tank during the day and night (Figure 4e-f), whereas *M. faiaziberi* stopped relatively more frequently and for more time, at the top of the tank during the day and toward the bottom at night (Figure 4g-h). Thus, *M.* sp. “Daktari”’s day-time tank usage involved near constant swimming along the bottom of the tank, while *M. faiaziberi’s* usage of the top of the tank involved more time stopping. Notable day/night differences were observed between these two species as well. Whereas *M. sp.* “Daktari” exhibited largely the same pattern during the day and night, *M. faiaziberi* showed a clear difference between day and night, with nearly all stops occurring toward the top of the tank during the day and toward the bottom at night.

## DISCUSSION

### Computer vision provides novel insights into cichlid behaviors

We provide a proof-of-concept for the biological relevance of our data produced through these CV methods. Further, these data extend existing inquiries into the roles of habitat partitioning in maintaining Lake Malawi cichlid biodiversity (Ford et al. 2016; Lloyd et al. 2024). We acknowledge that animal behavior in the lab may be markedly different from that in the wild; however, disentangling the myriad factors contributing to complex behaviors can be nearly impossible through field observations alone. On the other hand, laboratory studies allow for more precise control over environmental metrics, but these settings are necessarily artificial and may confound natural behaviors in several ways. Thus, lab and field studies complement one another, allowing a better understanding of animal behavior and its ecological relevance (List 2007; Aziz 2017). CV methods are well-suited to facilitate the cross-comparison of studies like these by revealing robust quantitative trends that would be difficult to glean using traditional methods. For instance, when manually scoring videos of animal behavior, one can draw conclusions about where an animal prefers to spend time, but such approaches are susceptible to user bias, error, and inconsistencies. Not only is CV more consistent, but it enables the acquisition and analysis of massive amounts of data to draw more robust conclusions about more subtle trends. Whether from the field or lab, these data can be used to generate hypotheses to be tested in the opposite realm. For example, not only did the behavioral patterns we characterized here aligned with these species’ known ecology, they also offered potential new insights into their behaviors. *A. labrosus* exhibited a strong spatial preference for tank corners, which aligns with their natural preference for rocky crevices in their natural environment (Oliver and Arnegard 2010). The same was true for *T. kumwera*, which barely stopped moving and appeared to have little spatial preference, as they are known menaces in the wild, vigorously guarding their particular patches of algae (Li et al., 2016). These were largely confirmatory data; however, our results also revealed new layers of understanding with respect to temporal niche partitioning. Specifically, *A. labrosus* exhibited a strong day-and-night pattern, whereby they occupied more tank space and stopped less at night. Nocturnal patterns of behavior have only recently been acknowledged in the conversation of niche partitioning in Malawi cichlids (Lloyd et al., 2021), and not yet for this species. These data raise the interesting possibility that *A. labrosus* may be nocturnal in the wild, a hypothesis that requires additional testing in the lab and field. In contrast to *A. labrosus* and *T. kumwera*, we had no a priori predictions regarding usage/stop patterns in either *Maylandia* species. These congeneric species possess similar ecomorphologies and accordingly exhibit largely similar diets and habitat preferences in the wild (Cicotto et al., 2011). Notably, our data revealed clear differences between tank usage and overall stops. In terms of usage, *M. sp*. “Daktari” exhibited a slight preference for the bottom of the tank during the day, while *M. faiaziberi* exhibited a strong preference for the top of the tank during the day. Additionally, *M. sp*. “Daktari” exhibited fewer stops over the entire twenty-four-hour filming period than *M. faiaziberi*. How or whether these patterns represent habitat partitioning in the wild remains an open question, but they suggest the possibility of partitioning via cryptic behaviors that would be difficult to observe in the wild. It is worth noting that these species were never co-housed in the lab. Thus, the behavioral variation noted here is unlikely due to each species segregating its movement patterns over time to accommodate the other. How mixing cichlid species with different behavioral phenotypes may influence individual behaviors is another interesting but open question. Overall, we feel that these types of analyses, even in a lab setting, can help to hone predictions about long-standing questions about how species navigate and partition their environment in space and time.

### Scalability of computer vision tools

Our analysis took large amounts of video data and reduced each fish to x, y coordinates that were captured every fifteenth of a second, leading to millions of data points collected over a twenty-four-hour period and a highly resolved view of which areas of the tank were routinely used and which were avoided. While the amount of data is considerable, the simplicity of each data point reduces the computational expense of the analysis. Furthermore, given the nature of the data (i.e., x, y), there were myriad ways to analyze/visualize them, providing a layered and nuanced understanding of animal activity. Specifically, our tank usage maps showed where fish existed in space but did not consider time. By adding the amount of time these animals spent at rest, we could speak to whether areas of high usage represented high-traffic “pass-through” regions of the tank or places where animals tended to stop/rest.

Another advantage of CV tools, in general, is that they may be scaled to produce much larger datasets than shown here; however, it bears mentioning that time spent training/retraining also scales with sample size, albeit not linearly. Iterative training is key to many CV projects as most models still struggle with generalization across large datasets due to subtle variability among samples. For instance, retraining may be required for videos of overall lower quality, fish with different color patterns, or when water is more turbid. This phenomenon is recognized as a problem in need of solving by computer scientists. For now, when generating larger datasets, biologists should be mindful of reducing variability within their control (e.g., water quality) and be willing to spend more time retraining their model for “outlier” videos. In spite of these considerations, the high-throughput potential of CV pairs very well with established biological methods. The search for the genetic basis and evolution of complex behaviors are two such lines of inquiry. Traditionally, these approaches require large sample sizes to achieve the necessary statistical power to draw meaningful conclusions. As a result, they are typically applied to anatomical phenotypes, especially in vertebrates. CV tools expand the ability to produce behavioral datasets with sample sizes sufficient for such analyses.

Understanding the neurogenetic mechanisms that underlie complex behaviors is an important topic that is well-studied in traditional laboratory models via genetic screens (Jain et al., 2019; Meserve et al., 2021), reverse genetics (Matynia et al., 2008; Howarth et al., 2023), or both (Takahashi et al., 2013). Meanwhile, the genetic mechanisms underlying variation or switches in these behaviors remain markedly understudied. A recent meta-analysis (Bubac et al., 2020) surveyed the literature for studies on the genetic basis of behavior in natural populations, and showed that not only were the overall numbers low (e.g., generally less than 20 per year since 2015), but about half took a candidate gene approach. Unbiased approaches such as GWAS and QTL mapping collectively accounted for the other half. The difficulty in collecting and the “messy” nature of behavioral data are likely root causes of the paucity of such studies. We suggest that CV methods offer a promising new tool in this realm.

The evolution of behavior is another ripe area of study that is benefiting from recent technological advances, both hardware and software (Hernández et al. 2021; Patricelli, 2023; Couzin and Heinz, 2023). For instance, recent work among insects underscores the potential for analyzing complex behaviors in a phylogenetic context (Raffudin and Crozier 2007; Hernández et al. 2021; York et al. 2022), but vertebrates again pose the logistical challenge of collecting sufficient amounts of data to draw meaningful conclusions about the evolution of complex behaviors. Notably, cichlids have proven to be a robust model for behavioral evolution. The building of elaborate and species-specific bowers by courting male cichlids, for example, has been examined in a phylogenetic context (Kidd et al., 2006), and associated with species-specific neuroanatomy and gene expression (York et al., 2019). Further, their experimental tractability in the lab has allowed for the recent application of CV techniques in classifying complex cichlid behaviors (Johnson et al. 2020; Lloyd et al. 2021; Lancaster et al. 2024). Thus, our pipeline adds to this incipient body of literature on the application of CV in cichlid behavior, with the potential to enhance an understanding of their molecular origins and evolution.

### Broader impacts of computer vision and artificial intelligence in science and society

We are watching a rapid rise in the use of all forms of Artificial Intelligence (AI), and as with every new technology, its application has pros and cons. Here, we show and discuss how a CV tool can increase the pace and consistency of data collection in a biological context. Though, even as we laude the uses of these tools, we must also acknowledge their downsides. While the broader societal considerations of AI are outside the scope of any scientific paper, there is plenty for the scientific community to consider as these technologies evolve and become more accessible. Some are already pushing back on the ‘AI revolution’, cautioning scientists to remain mindful of the application of these methods (Messeri and Crockett, 2024). We are inclined to agree with this perspective; despite our use of AI-derived tools, we see more space for amplifying current experimental designs rather than replacing them. True understanding of a small amount of data will always supersede misunderstanding a large volume of data. In an ideal world, all applications of AI tools would produce interpretable and relevant data, but there are limitations to both the technology and our own umwelt. Just as any non-scientific use of these tools should be inspected thoroughly, the application within our most trusted sources of information should be examined even more so. We expect and look forward to the scientific community considering and reflecting upon these technologies as they evolve and become more accessible and advance our chosen disciplines in dramatic and exciting ways.

## Acknowledgements

We thank everyone involved with the 2023 Resnick Sustainability Institute Summer Workshop on Computer Vision Methods for Ecology (CV4E Workshop). A special thanks to Tarun Sharma and Shir Bar for coding assistance and troubleshooting. Thanks also to members of the Albertson lab and the larger OEB community for feedback at multiple stages of this project. This research was funded by NSF/IOS 2128729 (to R.C.A and A.C.K), as well as the University of Massachusetts Graduate School Spaulding-Smith Fellowship (to N.H.).

## Data Availability

All data and code are available at https://github.com/QueenKrokuz/FD

## References

Adams, C.E., Huntingford, F.A. The functional significance of inherited differences in feeding morphology in a sympatric polymorphic population of Arctic charr. Evolutionary Ecology 16, 15–25 (2002). 10.1023/A:1016014124038

Albertson, R. & Pauers, Michael. (2019). “Morphological disparity in ecologically diverse versus constrained lineages of Lake Malaŵi rock-dwelling cichlids.” Hydrobiologia. 832. 10.1007/s10750-018-3829-z.

Alksne, Michaela N., et al. “Biogeographic Patterns of Pacific White-sided Dolphins Based on Long-term Passive Acoustic Records.” Diversity and Distributions, vol. 30, no. 9, Sept. 2024, p. e13903. DOI.org (Crossref), 10.1111/ddi.13903.

Arbour, J. H., and H. López-Fernández. “Adaptive Landscape and Functional Diversity of Neotropical Cichlids: Implications for the Ecology and Evolution of Cichlinae (Cichlidae; Cichliformes).” Journal of Evolutionary Biology, vol. 27, no. 11, Nov. 2014, pp. 2431–42. DOI.org (Crossref), 10.1111/jeb.12486.

Ardekani, Reza. “Computer Vision Approaches to the Analysis of Animal Behavior.” Order No. 3598193 University of Southern California, 2013. United States --California: ProQuest. Web. 4 Apr. 2025.

Aziz, Hassan. (2017). “Comparison between Field Research and Controlled Laboratory Research.” Archives of Clinical and Biomedical Research. 1. 101–104. 10.26502/acbr.50170011.

Baker, M. 1,500 scientists lift the lid on reproducibility. Nature 533, 452–454 (2016). 10.1038/533452a

Banerjee, Sreya, et al. “An Assistive Computer Vision Tool to Automatically Detect Changes in Fish Behavior in Response to Ambient Odor.” Scientific Reports, vol. 11, no. 1, Jan. 2021, p. 1002. *DOI.org (Crossref)*, 10.1038/s41598-020-79772-3.

Barnard, Shanis, et al. “Quick, Accurate, Smart: 3D Computer Vision Technology Helps Assessing Confined Animals’ Behaviour.” PLOS ONE, edited by Claire Wade, vol. 11, no. 7, July 2016, p. e0158748. *DOI.org (Crossref)*, 10.1371/journal.pone.0158748.

Bernardes, Rodrigo Cupertino, et al. “Ethoflow: Computer Vision and Artificial Intelligence-Based Software for Automatic Behavior Analysis.” Sensors, vol. 21, no. 9, May 2021, p. 3237. DOI.org (Crossref), 10.3390/s21093237.

Bewley, Alex, et al. “Simple Online and Realtime Tracking.” 2016 IEEE International Conference on Image Processing (ICIP), 2017, pp. 3464–68. arXiv.org, 10.1109/ICIP.2016.7533003.

Bhuiyan, Md Roman, and Philipp Wree. “Animal Behavior for Chicken Identification and Monitoring the Health Condition Using Computer Vision: A Systematic Review.” IEEE Access, vol. 11, 2023, pp. 126601–10. DOI.org (Crossref), 10.1109/ACCESS.2023.3331092.

Bootsma, Harvey A., et al. “Food Partitioning Among Lake Malawi Nearshore Fishes as Revealed by Stable Isotope Analyses.” Ecology, vol. 77, no. 4, June 1996, pp. 1286–90. DOI.org (Crossref), 10.2307/2265598.

Borges Oliveira, Dario Augusto, et al. “A Review of Deep Learning Algorithms for Computer Vision Systems in Livestock.” Livestock Science, vol. 253, Elsevier BV, Nov. 2021, p. 104700. Crossref, doi:10.1016/j.livsci.2021.104700.

Branch, Marc N. “The “Reproducibility Crisis:” Might the Methods Used Frequently in Behavior-Analysis Research Help?.” Perspectives on behavior science vol. 42,1 77–89. 4 Jun. 2018, doi:10.1007/s40614-018-0158-5

Broomé, Sofia, et al. “Going Deeper than Tracking: A Survey of Computer-Vision Based Recognition of Animal Pain and Affective States.” arXiv:2206.08405, arXiv, 16 June 2022. arXiv.org, 10.48550/arXiv.2206.08405.

Bubac, Christine M., et al. “The Genetic Basis of Animal Behavioural Diversity in Natural Populations.” Molecular Ecology, vol. 29, no. 11, June 2020, pp. 1957–71. DOI.org (Crossref), 10.1111/mec.15461.

Burress, Edward D. “Cichlid Fishes as Models of Ecological Diversification: Patterns, Mechanisms, and Consequences.” Hydrobiologia, vol. 748, no. 1, Apr. 2015, pp. 7–27. DOI.org (Crossref), 10.1007/s10750-014-1960-z.

Carruthers, Madeline, et al. “Ecological Speciation Promoted by Divergent Regulation of Functional Genes Within African Cichlid Fishes”, Molecular Biology and Evolution, Volume 39, Issue 11, November 2022, msac251, 10.1093/molbev/msac251

Chavez-Guerrero, Víctor Ocyel, et al. “Classification of Domestic Dogs Emotional Behavior Using Computer Vision.” Computación y Sistemas, vol. 26, no. 1, Mar. 2022. DOI.org (Crossref), 10.13053/cys-26-1-4165.

Chen, Chen, et al. “Behaviour Recognition of Pigs and Cattle: Journey from Computer Vision to Deep Learning.” Computers and Electronics in Agriculture, vol. 187, Aug. 2021, p. 106255. DOI.org (Crossref), 10.1016/j.compag.2021.106255.

Ciccotto, P.J., A. Konings and J.R. Stauffer Jr., 2011. “Descriptions of five new species in the genus Metriaclima (Teleostei: Cichlidae) from Lake Malaŵi”, Africa. Zootaxa 2738:1–25. (Ref. 86409)

Conith, Andrew J., et al. “Ecomorphological Divergence and Habitat Lability in the Context of Robust Patterns of Modularity in the Cichlid Feeding Apparatus.” BMC Evolutionary Biology, vol. 20, no. 1, Dec. 2020, p. 95. DOI.org (Crossref), 10.1186/s12862-020-01648-x.

Cooper, W. James, et al. “Bentho-Pelagic Divergence of Cichlid Feeding Architecture Was Prodigious and Consistent during Multiple Adaptive Radiations within African Rift-Lakes.” PLoS ONE, edited by Stuart Humphries, vol. 5, no. 3, Mar. 2010, p. e9551. *DOI.org (Crossref)*, 10.1371/journal.pone.0009551.

Couzin, Iain D., and Conor Heins. “Emerging Technologies for Behavioral Research in Changing Environments.” Trends in Ecology & Evolution, vol. 38, no. 4, Elsevier BV, Apr. 2023, pp. 346–354. Crossref, doi:10.1016/j.tree.2022.11.008.

Culjak, I., Abram, D., Pribanic, T., Dzapo, H., and Cifrek, M., “A brief introduction to OpenCV,” 2012 Proceedings of the 35th International Convention MIPRO, Opatija, Croatia, 2012, pp. 1725–1730.

CV4E Summer Workshop. https://cv4ecology.caltech.edu/.

Dave, Achal, et al. “Evaluating Large-Vocabulary Object Detectors: The Devil Is in the Details.” arXiv:2102.01066, arXiv, 15 Mar. 2022. arXiv.org, 10.48550/arXiv.2102.01066.

De León, L F et al. “Darwin’s finches and their diet niches: the sympatric coexistence of imperfect generalists.” Journal of evolutionary biology vol. 27,6 (2014): 1093–104. doi:10.1111/jeb.12383

Deng, J., Berg, A.C., Li, K., Fei-Fei, L. (2010). “What Does Classifying More Than 10,000 Image Categories Tell Us?.” In: Daniilidis, K., Maragos, P., Paragios, N. (eds) Computer Vision – ECCV 2010. ECCV 2010. Lecture Notes in Computer Science, vol 6315. Springer, Berlin, Heidelberg. 10.1007/978-3-642-15555-0_6

Doenz, Carmela J., et al. “Ecological Opportunity Shapes a Large Arctic Charr Species Radiation.” Proceedings of the Royal Society B: Biological Sciences, vol. 286, no. 1913, Oct. 2019, p. 20191992. DOI.org (Crossref), 10.1098/rspb.2019.1992.

Fan, J., Jiang, N., and Wu, Y., “Automatic video-based analysis of animal behaviors,” 2010 IEEE International Conference on Image Processing, Hong Kong, China, 2010, pp. 1513–1516, doi: 10.1109/ICIP.2010.5652495.

Fenton, Sam, et al. “Genomic Underpinnings of Head and Body Shape in Arctic Charr Ecomorph Pairs.” Molecular Ecology, vol. 33, no. 7, Apr. 2024, p. e17305. DOI.org (Crossref), 10.1111/mec.17305.

Fernandes, Arthur Francisco Araújo, et al. “Image Analysis and Computer Vision Applications in Animal Sciences: An Overview.” Frontiers in Veterinary Science, vol. 7, Oct. 2020, p. 551269. DOI.org (Crossref), 10.3389/fvets.2020.551269.

Ferrante, Gabriel S., et al. “Understanding the State of the Art in Animal Detection and Classification Using Computer Vision Technologies.” 2021 IEEE International Conference on Big Data (Big Data), IEEE, 2021, pp. 3056–65. DOI.org (Crossref), 10.1109/BigData52589.2021.9672049.

FFmpeg Developers. (2016). ffmpeg tool (Version be1d324) [Software]. Available from http://ffmpeg.org/

Flack, Andrea, et al. “From Local Collective Behavior to Global Migratory Patterns in White Storks.” Science, vol. 360, no. 6391, May 2018, pp. 911–14. DOI.org (Crossref), 10.1126/science.aap7781.

Ford, Antonia G. P., et al. “Niche Divergence Facilitated by Fine-scale Ecological Partitioning in a Recent Cichlid Fish Adaptive Radiation.” Evolution, vol. 70, no. 12, Dec. 2016, pp. 2718–35. DOI.org (Crossref), 10.1111/evo.13072.

Gavrilets, Sergey, et al. “Case Studies and Mathematical Models of Ecological Speciation. 1. Cichlids in a Crater Lake.” Molecular Ecology, vol. 16, no. 14, July 2007, pp. 2893–909. DOI.org (Crossref), 10.1111/j.1365-294X.2007.03305.x.

Genner, et al. “Niche Segregation among Lake Malawi Cichlid Fishes? Evidence from Stable Isotope Signatures.” Ecology Letters, vol. 2, no. 3, December. 2001, pp. 185–90. DOI.org (Crossref), 10.1046/j.1461-0248.1999.00068.x.

Genner, M. J., et al. “Age of Cichlids: New Dates for Ancient Lake Fish Radiations.” Molecular Biology and Evolution, vol. 24, no. 5, Feb. 2007, pp. 1269–82. DOI.org (Crossref), 10.1093/molbev/msm050.

Genner, Martin J et al. “Foraging of rocky habitat cichlid fishes in Lake Malawi: coexistence through niche partitioning?.” Oecologia vol. 121,2 (1999): 283–292. doi:10.1007/s004420050930

Gould, J. 1837. [Remarks on a Group of Ground Finches from Mr. Darwin’s Collection, with Characters of the New Species]. Proceedings of the Zoological Society of London 5: 4–7.

Graving, Jacob M. “Computer Vision and Deep Learning Methods for Measuring and Modeling Animal Behavior.”

Hassan, Aziz. “Comparison between Field Research and Controlled Laboratory Research.” Archives of Clinical and Biomedical Research, vol. 01, no. 02, 2017, pp. 101–04. DOI.org (Crossref), 10.26502/acbr.50170011.

Henderson, Paul, and Vittorio Ferrari. “End-to-End Training of Object Class Detectors for Mean Average Precision.” arXiv:1607.03476, arXiv, 16 Mar. 2017. arXiv.org, 10.48550/arXiv.1607.03476.

Hernández, Damián G., et al. “A Framework for Studying Behavioral Evolution by Reconstructing Ancestral Repertoires.” eLife, vol. 10, Sept. 2021, p. e61806. DOI.org (Crossref), 10.7554/eLife.61806.

Howarth, Emmeline R. I., et al. “Genetic Polymorphisms in the Serotonin, Dopamine and Opioid Pathways Influence Social Attention in Rhesus Macaques (Macaca Mulatta).” PLOS ONE, edited by Asem Surindro Singh, vol. 18, no. 8, Aug. 2023, p. e0288108. DOI.org (Crossref), 10.1371/journal.pone.0288108.

Hutchison, David, et al. “What Does Classifying More Than 10,000 Image Categories Tell Us?” Computer Vision – ECCV 2010, edited by Kostas Daniilidis et al., vol. 6315, Springer Berlin Heidelberg, 2010, pp. 71–84. *DOI.org (Crossref)*, 10.1007/978-3-642-15555-0_6.

Ingram, T., et al. “Hierarchical Partitioning of Multiple Niche Dimensions among Ecomorphs, Species and Sexes in Puerto Rican Anoles.” Journal of Zoology, vol. 318, no. 2, Wiley, 20 July 2022, pp. 127–134. Crossref, doi:10.1111/jzo.13004.

Jain, Puneet, et al. “Development of Criteria for Epilepsy Genetic Testing in Ontario, Canada.” Canadian Journal of Neurological Sciences / Journal Canadien Des Sciences Neurologiques, vol. 46, no. 1, Jan. 2019, pp. 7–13. DOI.org (Crossref), 10.1017/cjn.2018.341.

Jiang, Yunfan “CS231n Deep Learning for Computer Vision.” 2015. Github.Io., from https://cs231n.github.io/.

Johnson, Zachary V., et al. “Automated Measurement of Long-Term Bower Behaviors in Lake Malawi Cichlids Using Depth Sensing and Action Recognition.” Scientific Reports, vol. 10, no. 1, Nov. 2020, p. 20573. DOI.org (Crossref), 10.1038/s41598-020-77549-2.

Jordan, A., Taborsky, B., Taborsky, M. (2021). “Cichlids as a Model System for Studying Social Behaviour and Evolution.” In: Abate, M.E., Noakes, D.L. (eds) The Behavior, Ecology and Evolution of Cichlid Fishes. Fish & Fisheries Series, vol 40. Springer, Dordrecht. 10.1007/978-94-024-2080-7_16

Khanam, Rahima, and Muhammad Hussain. “What Is YOLOv5: A Deep Look into the Internal Features of the Popular Object Detector.” arXiv:2407.20892, arXiv, 30 July 2024. arXiv.org, 10.48550/arXiv.2407.20892.

Kidd, Michael R., et al. “Axes of Differentiation in the Bower-building Cichlids of Lake Malawi.” Molecular Ecology, vol. 15, no. 2, Feb. 2006, pp. 459–78. DOI.org (Crossref), 10.1111/j.1365-294X.2005.02787.x.

Koblmüller, Stephan, et al. “Preface: Advances in Cichlid Research III: Behavior, Ecology, and Evolutionary Biology.” Hydrobiologia, vol. 832, no. 1, Apr. 2019, pp. 1–8. DOI.org (Crossref), 10.1007/s10750-019-3903-1.

Kocher, Thomas D. “Adaptive Evolution and Explosive Speciation: The Cichlid Fish Model.” Nature Reviews Genetics, vol. 5, no. 4, Springer Science and Business Media LLC, 1 Apr. 2004, pp. 288–298. Crossref, doi:10.1038/nrg1316.

Koger, Benjamin, et al. “Quantifying the Movement, Behaviour and Environmental Context of Group-living Animals Using Drones and Computer Vision.” Journal of Animal Ecology, vol. 92, no. 7, July 2023, pp. 1357–71. DOI.org (Crossref), 10.1111/1365-2656.13904.

Kornfield, Irv, and Peter F. Smith. “African Cichlid Fishes: Model Systems for Evolutionary Biology.” Annual Review of Ecology and Systematics, vol. 31, no. 1, Nov. 2000, pp. 163–96. DOI.org (Crossref), 10.1146/annurev.ecolsys.31.1.163.

Kullander, S.O., 1998. “A phylogeny and classification of the South American Cichlidae (Teleostei: Perciformes).” p. 461-498. In L.R. Malabarba, R.E. Reis, R.P. Vari, Z.M. Lucena and C.A.S. Lucena (eds.) Phylogeny and classification of neotropical fishes. Porto Alegre, Edipucrs. 603 p.

Lancaster, Tucker J., et al. “SARTAB, a Scalable System for Automated Real-Time Behavior Detection Based on Animal Tracking and Region Of Interest Analysis: Validation on Fish Courtship Behavior.” Frontiers in Behavioral Neuroscience, vol. 18, Dec. 2024, p. 1509369. DOI.org (Crossref), 10.3389/fnbeh.2024.1509369.

Lao, F., et al. “Behavior recognition method for individual laying hen based on computer vision.” Transactions of the Chinese Society of Agricultural Engineering, Volume 28, Number 24, 15 December 2012, pp. 157–163(7)

Leroy, T., Vranken, E., Van Brecht, A., et al. “A COMPUTER VISION METHOD FOR ON-LINE BEHAVIORAL QUANTIFICATION OF INDIVIDUALLY CAGED POULTRY.” Transactions of the ASABE, vol. 49, no. 3, 2006, pp. 795–802. DOI.org (Crossref), 10.13031/2013.20462.

Leroy, T., Vranken, E., Struelens, E., et al. “Computer Vision Based Recognition of Behavior Phenotypes of Laying Hens.” 2005 Tampa, FL July 17-20, 2005, American Society of Agricultural and Biological Engineers, 2005. DOI.org (Crossref), 10.13031/2013.19471.

Li, Guoming, et al. “Practices and Applications of Convolutional Neural Network-Based Computer Vision Systems in Animal Farming: A Review.” Sensors, vol. 21, no. 4, Feb. 2021, p. 1492. DOI.org (Crossref), 10.3390/s21041492.

Li, S., A.F. Konings and J.R. Stauffer Jr., 2016. “A revision of the Pseudotropheus elongatus species group (Teleostei: Cichlidae) with description of a new genus and seven new species.” Zootaxa 4168(2):353–381. (Ref. 119465)

Lindeberg, T. Feature “Detection with Automatic Scale Selection.” International Journal of Computer Vision 30, 79–116 (1998). 10.1023/A:1008045108935

Liu, Dong, et al. “A Computer Vision-Based Method for Spatial-Temporal Action Recognition of Tail-Biting Behaviour in Group-Housed Pigs.” Biosystems Engineering, vol. 195, Elsevier BV, July 2020, pp. 27–41. Crossref, doi:10.1016/j.biosystemseng.2020.04.007.

List, John. “Field Experiments: A Bridge Between Lab and Naturally-Occurring Data. National Bureau of Economic Research”, Mar. 2007. Crossref, doi:10.3386/w12992.

Lloyd, Evan, et al. “Diversity in Rest–Activity Patterns among Lake Malawi Cichlid Fishes Suggests a Novel Axis of Habitat Partitioning.” Journal of Experimental Biology, vol. 224, no. 7, Apr. 2021, p. jeb242186. DOI.org (Crossref), 10.1242/jeb.242186.

Lloyd, Evan, et al. “Ontogeny and Social Context Regulate the Circadian Activity Patterns of Lake Malawi Cichlids.” Journal of Comparative Physiology B, vol. 194, no. 3, June 2024, pp. 299–313. DOI.org (Crossref), 10.1007/s00360-023-01523-3.

Lopez-Fuentes, Laura, et al. “Review on Computer Vision Techniques in Emergency Situations.” Multimedia Tools and Applications, vol. 77, no. 13, July 2018, pp. 17069–107. DOI.org (Crossref), 10.1007/s11042-017-5276-7.

Lowe, David G. “Distinctive Image Features from Scale-Invariant Keypoints.” International Journal of Computer Vision, vol. 60, no. 2, Nov. 2004, pp. 91–110. DOI.org (Crossref), 10.1023/B:VISI.0000029664.99615.94.

MacArthur, Robert H. “Population Ecology of Some Warblers of Northeastern Coniferous Forests.” Ecology, vol. 39, no. 4, Oct. 1958, pp. 599–619. DOI.org (Crossref), 10.2307/1931600.

Martin, Christopher H., and Martin J. Genner. “High Niche Overlap between Two Successfully Coexisting Pairs of Lake Malawi Cichlid Fishes.” Canadian Journal of Fisheries and Aquatic Sciences, vol. 66, no. 4, Apr. 2009, pp. 579–88. DOI.org (Crossref), 10.1139/F09-023.

Martinez, Christopher M., et al. “Feeding Ecology Underlies the Evolution of Cichlid Jaw Mobility.” Evolution, vol. 72, no. 8, Aug. 2018, pp. 1645–55. DOI.org (Crossref), 10.1111/evo.13518.

Mathis, Alexander, et al. “DeepLabCut: Markerless Pose Estimation of User-Defined Body Parts with Deep Learning.” Nature Neuroscience, vol. 21, no. 9, Sept. 2018, pp. 1281–89. DOI.org (Crossref), 10.1038/s41593-018-0209-y.

Mathis, Mackenzie Weygandt, and Alexander Mathis. “Deep Learning Tools for the Measurement of Animal Behavior in Neuroscience.” Current Opinion in Neurobiology, vol. 60, Feb. 2020, pp. 1–11. DOI.org (Crossref), 10.1016/j.conb.2019.10.008.

Matynia, Anna, et al. “A High Through-Put Reverse Genetic Screen Identifies Two Genes Involved in Remote Memory in Mice.” PLoS ONE, edited by Wim E. Crusio, vol. 3, no. 5, May 2008, p. e2121. DOI.org (Crossref), 10.1371/journal.pone.0002121.

Meserve, Joy H., et al. “A Forward Genetic Screen Identifies Dolk as a Regulator of Startle Magnitude through the Potassium Channel Subunit Kv1.1.” PLOS Genetics, edited by Cecilia Moens, vol. 17, no. 6, June 2021, p. e1008943. DOI.org (Crossref), 10.1371/journal.pgen.1008943.

Messeri, Lisa, and M. J. Crockett. “Artificial Intelligence and Illusions of Understanding in Scientific Research.” Nature, vol. 627, no. 8002, Mar. 2024, pp. 49–58. DOI.org

Mönck, Hauke Jürgen, et al. “BioTracker: An Open-Source Computer Vision Framework for Visual Animal Tracking.” arXiv:1803.07985, arXiv, 21 Mar. 2018. arXiv.org, 10.48550/arXiv.1803.07985.

Nichols, Annika L. A., et al. “Widespread Temporal Niche Partitioning in an Adaptive Radiation of Cichlid Fishes.” Evolutionary Biology, 2 June 2024. DOI.org (Crossref), 10.1101/2024.05.29.596472.

Noldus, Lucas P. J. J., et al. “EthoVision: A Versatile Video Tracking System for Automation of Behavioral Experiments.” Behavior Research Methods, Instruments, & Computers, vol. 33, no. 3, Aug. 2001, pp. 398–414. DOI.org (Crossref), 10.3758/BF03195394.

Oliver, M.K. and M.E. Arnegard, 2010. “A new genus for Melanochromis labrosus, a problematic Lake Malawi cichlid with hypertrophied lips (Teleostei: Cichlidae).” Ichthyol. Explor. Freshwat. 21(3):209–232. (Ref. 85491)

Okuyama, Junichi, et al. “Application of a Computer Vision Technique to Animal-Borne Video Data: Extraction of Head Movement to Understand Sea Turtles’ Visual Assessment of Surroundings.” Animal Biotelemetry, vol. 3, no. 1, Dec. 2015, p. 35. DOI.org (Crossref), 10.1186/s40317-015-0079-y.

Oyen, Diane, et al. “Controlling the Precision-Recall Tradeoff in Differential Dependency Network Analysis.” arXiv:1307.2611, arXiv, 9 July 2013. arXiv.org, 10.48550/arXiv.1307.2611.

Palmér, Tobias. “Computer Vision Based Analysis of Animal Behavior.” Mathematics, Centre for Mathematical Sciences, Faculty of Engineering, Lund University, 2017.

Papadakis, Vassilis M., et al. “A Computer-Vision System and Methodology for the Analysis of Fish Behavior.” Aquacultural Engineering, vol. 46, Jan. 2012, pp. 53–59. DOI.org (Crossref), 10.1016/j.aquaeng.2011.11.002.

Parisot, Sarah, et al. “Learning to Name Classes for Vision and Language Models.” arXiv:2304.01830, arXiv, 4 Apr. 2023. arXiv.org, 10.48550/arXiv.2304.01830.

Parsons, Kevin J., Skuli Skúlason, et al. “Morphological Variation over Ontogeny and Environments in Resource Polymorphic Arctic Charr (Salvelinus Alpinus).” Evolution & Development, vol. 12, no. 3, May 2010, pp. 246–57. DOI.org (Crossref), 10.1111/j.1525-142X.2010.00410.x.

Parsons, Kevin J., Moira Concannon, et al. “Foraging Environment Determines the Genetic Architecture and Evolutionary Potential of Trophic Morphology in Cichlid Fishes.” Molecular Ecology, vol. 25, no. 24, Dec. 2016, pp. 6012–23. DOI.org (Crossref), 10.1111/mec.13801.

Patricelli GL (2023) Behavioral ecology: New technology enables a more holistic view of complex animal behavior. PLoS Biol 21(8): e3002264. 10.1371/journal.pbio.3002264

Pedregosa, Fabian, et al. “Scikit-Learn: Machine Learning in Python.” arXiv:1201.0490, arXiv, 5 June 2018. arXiv.org, 10.48550/arXiv.1201.0490.

Pereira, Talmo D., et al. “SLEAP: A Deep Learning System for Multi-Animal Pose Tracking.” Nature Methods, vol. 19, no. 4, Apr. 2022, pp. 486–95. DOI.org (Crossref), 10.1038/s41592-022-01426-1.

Raffiudin, Rika, and Ross H. Crozier. “Phylogenetic Analysis of Honey Bee Behavioral Evolution.” Molecular Phylogenetics and Evolution, vol. 43, no. 2, May 2007, pp. 543–52. DOI.org (Crossref), 10.1016/j.ympev.2006.10.013.

Redmon, Joseph, et al. “You Only Look Once: Unified, Real-Time Object Detection.” arXiv:1506.02640, arXiv, 9 May 2016. arXiv.org, 10.48550/arXiv.1506.02640.

Ribbink, A. J., et al. “A Preliminary Survey of the Cichlid Fishes of Rocky Habitats in Lake Malawi.” South African Journal of Zoology, vol. 18, no. 3, Jan. 1983, pp. 149–310. DOI.org (Crossref), 10.1080/02541858.1983.11447831.

Santos, M.E., Lopes, J.F. & Kratochwil, C.F. East African cichlid fishes. EvoDevo 14, 1 (2023). 10.1186/s13227-022-00205-5

Sampaio, Ana Lúcia A., et al. “Relationships between Morphology, Diet and Spatial Distribution: Testing the Effects of Intra and Interspecific Morphological Variations on the Patterns of Resource Use in Two Neotropical Cichlids.” Neotropical Ichthyology, vol. 11, no. 2, June 2013, pp. 351–60. DOI.org (Crossref), 10.1590/S1679-62252013005000001.

Schneider, Stefan, et al. “Past, Present and Future Approaches Using Computer Vision for Animal Re-identification from Camera Trap Data.” Methods in Ecology and Evolution, edited by Robert B. O’Hara, vol. 10, no. 4, December. 2018, pp. 461–70. *DOI.org (Crossref)*, 10.1111/2041-210X.13133.

Schütz, Anne K., et al. “Computer Vision for Detection of Body Posture and Behavior of Red Foxes.” Animals, vol. 12, no. 3, Jan. 2022, p. 233. DOI.org (Crossref), 10.3390/ani12030233.

Seehausen, Ole. “African Cichlid Fish: A Model System in Adaptive Radiation Research.” Proceedings of the Royal Society B: Biological Sciences, vol. 273, no. 1597, Aug. 2006, pp. 1987–98. DOI.org (Crossref), 10.1098/rspb.2006.3539.

Selfridge, Oliver “Pandemonium: A paradigm for learning” Symposium on the mechanization of thought processes, 1959, HM Stationary Office, London

Skoglund, Sigrid, et al. “Morphological Divergence between Three Arctic Charr Morphs – the Significance of the Deep-water Environment.” Ecology and Evolution, vol. 5, no. 15, Wiley, 14 July 2015, pp. 3114–3129. Crossref, doi:10.1002/ece3.1573.

Solsona-Berga, Alba, et al. “Machine Learning with Taxonomic Family Delimitation Aids in the Classification of Ephemeral Beaked Whale Events in Passive Acoustic Monitoring.” PLOS ONE, edited by Steffen Kiel, vol. 19, no. 6, June 2024, p. e0304744. DOI.org (Crossref), 10.1371/journal.pone.0304744.

Soria-Barreto, Miriam, and Rocío Rodiles-Hernández. “Spatial Distribution of Cichlids in Tzendales River, Biosphere Reserve Montes Azules, Chiapas, Mexico.” Environmental Biology of Fishes, vol. 83, no. 4, Dec. 2008, pp. 459–69. DOI.org (Crossref), 10.1007/s10641-008-9368-0.

Stefanics, Daniela, and Markus Fox. “COCO Annotator.” ACM SIGMultimedia Records, vol. 13, no. 3, Association for Computing Machinery (ACM), December. 2022, pp. 1–1. Crossref, doi:10.1145/3578495.3578502.

Stern, Ulrich, et al. “Analyzing Animal Behavior via Classifying Each Video Frame Using Convolutional Neural Networks.” Scientific Reports, vol. 5, no. 1, Sept. 2015, p. 14351. DOI.org (Crossref), 10.1038/srep14351.

Takahashi, Joseph S., et al. “Forward and Reverse Genetic Approaches to Behavior in the Mouse.” Science, vol. 264, no. 5166, June 1994, pp. 1724–33. DOI.org (Crossref), 10.1126/science.8209253.

Tetrault, Emily, Ben Aaronson, et al. “Foraging-Induced Craniofacial Plasticity Is Associated with an Early, Robust and Dynamic Transcriptional Response.” Proceedings of the Royal Society B: Biological Sciences, vol. 291, no. 2021, Apr. 2024, p. 20240215. DOI.org (Crossref), 10.1098/rspb.2024.0215.

Tkachenko, Maxim et al., 2020. Label Studio: Data labeling software, Available at: https://github.com/heartexlabs/label-studio.

Trejo, Karla, and Cecilio Angulo. “Single-Camera Automatic Landmarking for People Recognition with an Ensemble of Regression Trees.” Computación y Sistemas, vol. 20, no. 1, Mar. 2016. DOI.org (Crossref), 10.13053/cys-20-1-2365.

Vogg, Richard, et al. Computer Vision for Primate Behavior Analysis in the Wild. arXiv:2401.16424, arXiv, 12 Aug. 2024. arXiv.org, 10.48550/arXiv.2401.16424.

Walter, Tristan Leonard. “Efficient Computer Vision and Machine Learning Methods for Automating Large-Scale Analysis of Collective Animal Behavior.”

Wijeyakulasuriya, Dhanushi A., et al. “Machine Learning for Modeling Animal Movement.” PLOS ONE, edited by Jie Zhang, vol. 15, no. 7, July 2020, p. e0235750. DOI.org (Crossref), 10.1371/journal.pone.0235750.

Wiley, Victor, and Thomas Lucas. “Computer Vision and Image Processing: A Paper Review.” International Journal of Artificial Intelligence Research, vol. 2, no. 1, June 2018, p. 22. DOI.org (Crossref), 10.29099/ijair.v2i1.42.

Wurtz, Kaitlin, et al. “Recording Behaviour of Indoor-Housed Farm Animals Automatically Using Machine Vision Technology: A Systematic Review.” PLOS ONE, edited by Didier Raboisson, vol. 14, no. 12, Dec. 2019, p. e0226669. *DOI.org (Crossref)*, 10.1371/journal.pone.0226669.

Yilmaz, Alper, et al. “Object Tracking: A Survey.” ACM Computing Surveys, vol. 38, no. 4, Dec. 2006, p. 13. DOI.org (Crossref), 10.1145/1177352.1177355.

York, Ryan A., Allie Byrne, et al. “Behavioral Evolution Contributes to Hindbrain Diversification among Lake Malawi Cichlid Fish.” Scientific Reports, vol. 9, no. 1, Dec. 2019, p. 19994. DOI.org (Crossref), 10.1038/s41598-019-55894-1.

York, Ryan A., Luke E. Brezovec, et al. “The Evolutionary Trajectory of Drosophilid Walking.” Current Biology, vol. 32, no. 14, July 2022, pp. 3005–3015.e6. DOI.org (Crossref), 10.1016/j.cub.2022.05.039.

Zhao, Xia, et al. “A Review of Convolutional Neural Networks in Computer Vision.” Artificial Intelligence Review, vol. 57, no. 4, Mar. 2024, p. 99. DOI.org (Crossref), 10.1007/s10462-024-10721-6.

